# Thermo- and photoreceptors of eyes as regulators of circadian rhythm and glymphatic system

**DOI:** 10.1101/2023.09.15.557924

**Authors:** Alexander Kholmanskiy

**Affiliations:** Science Center Bemcom, Moscow, Russia

**Keywords:** temperature, water, eyes, brain, circadian rhythm, glymphatic system

## Abstract

The dynamics of hydrogen bonds in bulk and hydrated water affected the activation energies of temperature dependence of ion currents of voltage-dependent channels that regulate communication and trophic bonds in the neuropil of the cortical parenchyma. The physics of minimizing of isobaric heat capacity of water made it possible to explain stabilization and functional optimization of thermodynamics of eyeball fluids at 34.5 °C and human brain during sleep at 36.5 °C. At these temperatures, thermoreceptors of cornea and cells of ganglionic layer of the retina, through connections with suprachiasmatic nucleus and pineal gland, switch brain metabolism from daytime to nighttime modes. The phylogenesis of circadian rhythm was reflected in dependence of duration of nighttime sleep of mammals on diameter of eyeball, mass of pineal gland, and density of neurons in parenchyma of cortex. The activity of all nerves of eyeball led to division of nocturnal sleep into slow and fast phases. These phases correspond to two modes of glymphatic system - electrochemical and dynamic. The first is responsible for relaxation processes of synaptic plasticity and chemical neutralization of toxins with participation of water and melatonin. Rapid eye movement and an increase in cerebral blood flow in second mode increase water exchange in parenchyma and flush out toxins into venous system.

## 1. Introduction

The global biogenic factors of the circadian rhythm are sunlight during the day and cold at night. Their influence caused the emergence and development of receptors in animals that react to visible light and cold. In mammals, these receptors are localized in the eyeball in combination with the vitreous body (VB), which consists of ∼99% water. The ganglionic layer of the photosensitive retina is adjacent to the dorsal surface of the VB, and the thermoreceptors are concentrated in the cornea, which has close thermal contact with the VB through the lens and aqueous humor of the eye chambers, which is similar in composition and functions to blood plasma. The eyeball is thermally isolated from the bones of the orbit by a layer of adipose tissue and from the external environment by eyelids, the fiber of which is devoid of a fatty layer. Moreover, the average temperature (T) of the cornea and VB, equal to 34.5±0.5 °C [1], coincides with T_w_, in which the temperature dependence (TD) of the isobaric heat capacity of water (C_P_) has a flat minimum [2, 3]. The thermodynamic features of water in the vicinity of T_w_ and the thermal insulation of the brain by the cranial bone and scalp provide stabilization of human brain metabolism during sleep at T_S_=36.5 °C and during wakefulness T_b_∼37 °C [4-6].

The gel-like substance VB within its own fibrillar sheath is «reinforced» with threads of collagen and hyaluronic acid. During the transition from wakefulness to sleep, the flow rate of aqueous humor along VB decreases by almost half [1]. At a qualitative level, the circadian rhythm manifests itself in intraocular pressure [7], and in the rapid eye movement sleep phase (REM phase) pressure fluctuations were minimal, and at various stages of slow-wave sleep (NREM phase) they reached a maximum in spindle oscillations [8, 9]. These fluctuations will modulate the movement of ocular fluid along the VB into the perivascular space of the optic nerve [10], simultaneously participating, together with capillary water, in the cleansing drainage of the retina in a state of sleep or drowsiness. The density of nerve endings of the trigeminal nerve that respond to heat (pain) and cold in the human cornea is two orders of magnitude higher than in the skin of the fingers [11-14]. The high sensitivity of corneal cold receptors is due to their membrane voltage-gated cation channels TRPM8 [15, 16]. Cells of the ganglion layer complete the transformation of light information into optic nerve impulses, which, after primary processing in the thalamus, are fan-fed to the visual area of the cerebral cortex [17]. At the same time, in the ganglion layer, ∼1-2% of cells (ipRGC) are classified as special – they synchronize their physiology and activity with the circadian «light-dark» cycle [18]. ipRGC cells have their own photopigment, melanopsin, different from rhodopsin, and are directly connected to the suprachiasmatic nucleus (SCN) of the hypothalamus [19–22].

For animals and humans, here are established correlations between sleep duration and anatomical parameters of the eyeball, pineal gland and cerebral cortex. Analysis of the known dependences of neural activity of the eyes on temperature confirmed the participation of special cells of the ganglion layer of the retina and cold receptors of the cornea in the mechanism of switching brain metabolism from day to night mode and rapid sleep to slow sleep.

## 2. Methods and materials

The TD molecular dynamics features of pure water are due to a combination of Brownian motion of individual molecules and collective librational vibrations of molecules in a network of hydrogen bonds (HBs). Effective activation energies (EA), obtained from Arrhenius approximations of TDs parameters of rotational-translational dynamics of molecules, are determined by the energies of breaking hydrogen bonds and exit of molecule from the cell. The average number of tetrahedral HBs per water molecule in the range of 25-40 °C is ∼3-3.5 [23], therefore the moduli of E_A_ values for the self-diffusion coefficient (D_w_), viscosity, spin (T_1_) and dielectric relaxation (τ_D_) vary in the range from ∼15 to ∼20 kJ/mol with thermal energy (E_T_) of the order of ∼2.5 kJ/mol [2, 24]. On the other hand, the E_A_ of cooperative transformations in the HBs network will be determined by the balance of the energy of thermal fluctuations of the HBs angles (∼E_T_) and the electrical energy (E_R_) of proton hopping inside the cell and dipole-dipole interactions between molecules [3, 25-27].

In the general case, taking into account the known forms of the dependence of the properties of water on T^β^, the contribution of E_T_ in the Arrhenius approximations (F_A_) was expressed as a separate exponential [25, 28, 29]:

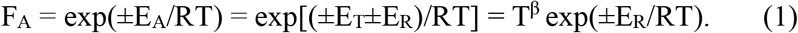

R – gas constant (8.31 J·mol-1·K-1). The plus and minus signs correspond to the thermal effect of exo and endothermic restructuring of HBs. Application of (1) made it possible to obtain reliable E_A_ values for all water parameters with known TDs and explain the nature of TDs anomalies. For example, for C_P_ β=1, and for water density and dielectric constant β=-1 [30].

The dynamics of bulk water and metabolites in blood and cerebrospinal fluid (CSF) *in vivo* are modulated by arterial and osmotic pressure gradients. The mobility of dipoles of water, ions and charged metabolites at the microlevel and in transmembrane channels is stimulated by gradients of local electric fields of polarized media and charged surfaces. The dynamics and transformations of proteins in physiological fluids *in vivo* and in model solutions *in vito* are limited by the transformations of HBs in the hydration shells of proteins, the total energy of which can be significantly more than 20 kJ/mol. Accordingly, E_A_ for TDs of dynamic, electrical and optical properties of model protein solutions vary from ∼50 to ∼400 kJ/mol [30-32]. These data were taken into account when analyzing known TDs the properties of such solutions using (1). If necessary, TDs were divided into T-intervals, in which the E_A_ values in (1) corresponded to the dominant factor of solution dynamics. The reliability of approximations (1) and E_A_ values was determined by the degree of closeness of R^2^ to 1.

## 3. Results and discussion

### 3.1. Physiological correlations

In mammals, when falling asleep, ipRGC cells activate the SCN and its output neurohumoral signals switch the metabolic mode of the pineal gland [6, 22, 33-35]. Increases in the blood and CSF, the content of melatonin, serotonin, norepinephrine and other neurotransmitters, provide switching the homeostasis of the cerebral cortex and hydrodynamics of CSF to the glymphatic system mode. Its productivity is determined by the physiology of sleep and is proportional to the density of neurons in the neocortex, since the level of contamination of the intercellular fluid (ICF) of the parenchyma depends on their total activity during wakefulness. Genetically, the need for neocortical clearance reflects the specific lifestyle and physical characteristics of the animal’s habitat. These features in mammals should correlate with the physiological parameters of sleep and the circadian rhythm control system, the signal chain of which includes the eyes with their photo- and thermoreceptors, the SCN, the pineal gland and the sleep center in the midbrain [18, 22].

Genetic features of the circadian rhythm control system in mammals were manifested on the pineal gland, which is absent in the electric ray, crocodile, and cetaceans, not found in the dolphin, and very small in the elephant [36]. In mammals of northern latitudes, the pineal gland is larger than in inhabitants of southern latitudes. Average volume of the pineal gland (in mm^3^): 6÷12 (predators); 60÷300 (ungulates); ∼180 (monkey); ∼200 (person) [36]. A similar correlation among terrestrial mammals is observed in the form of a direct dependence of sleep duration on the diameter of their eyeball (d_e_) (Fig. 1). The value of d_e_ determines the volume of VB, the area of the cornea and the number of ipRGC cells in the ganglion layer of the retina. A clear confirmation of these correlations is ∼2 hours of sleep-in elephants, which have twice the cortical mass of humans, but 3 times less neuron density compared to the human cortex [37]. The absence of the pineal gland and the REM phase in dolphins and cetaceans [38] is apparently due to the limitation of daily fluctuations in the temperature of the world ocean by ∼0.3 °C [39] and the possible leveling by the aquatic environment of the difference in the effect of the biogenic solar factor on the biosphere day and night [33, 40].

**Fig. 1.**
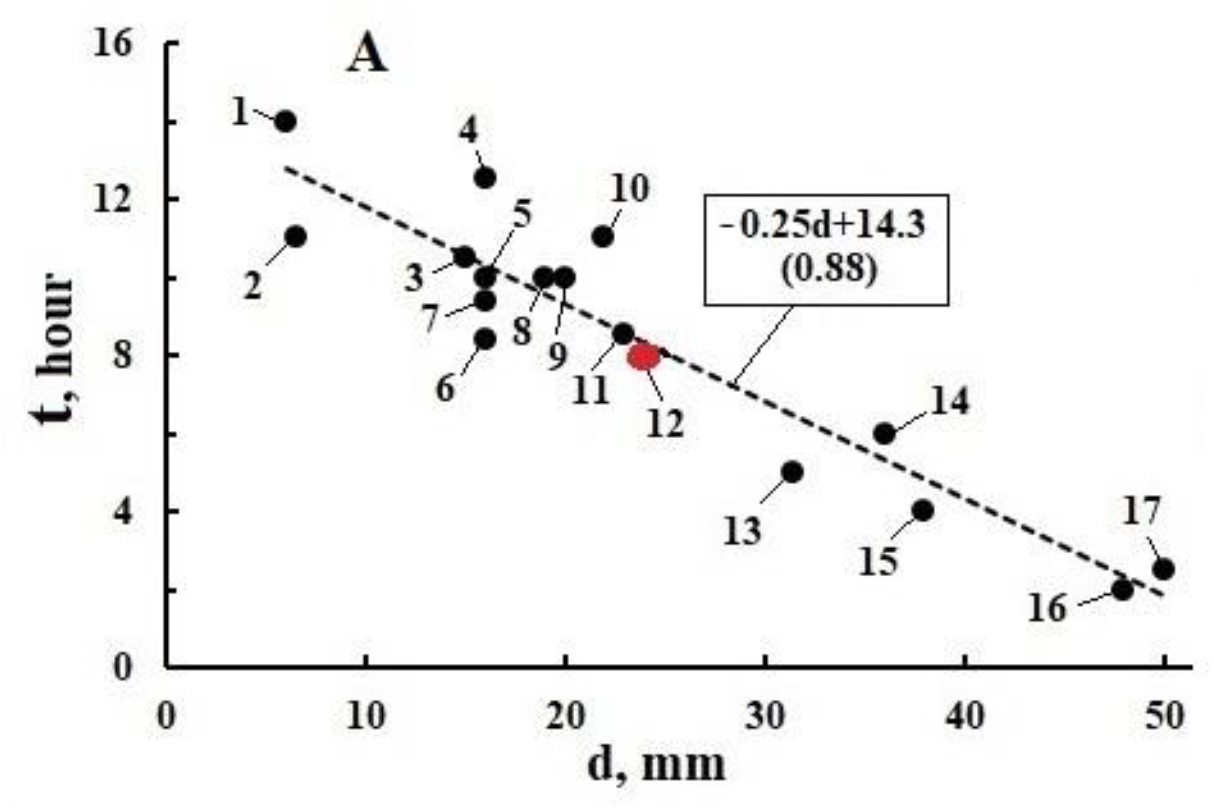
Dependence of the duration of sleep per day (t) of mammals on the diameter of their eyeball (d_e_): 1 - platypus, 2 - rat, 3 - adult squirrel, 4 - cat, 5 - fox, 6 - rabbit, 7 - lemur mouse, 8 - rhesus monkey, 9 - dog, 10 - orangutan, 11 - pig, 12 - Human, 13 - goat, 14 - sea lion, 15 - elephant, 16 - giraffe, 17 - horse. Open data on GOOGLE.

### 3.2. Two modes of the glymphatic system

A normal electroencephalogram in a state of sleep is divided into phases NREM and REM [41-44], the physiology of which corresponds to two conditional modes of operation of the glymphatic system – electrochemical (GS1) and dynamic (GS2) [6, 45, 46]. When falling asleep in the GS1 mode, different stages of NREM sleep dominate and the eye muscles and neuropil are periodically fed with glucose for 15-20 minutes [47-49]. At the same time, in the parenchyma the level of glucose and oxygen consumption does not change significantly [50, 51]. In the GS1 mode, relaxation processes of synaptic plasticity and chemical neutralization of toxins occur with the participation of water and melatonin [52-54]. In GS1, a decrease in cerebral blood flow by ∼25% and blood volume by ∼10% is accompanied by an influx of CSF into the third and fourth ventricles [56, 57]. It can be assumed that a decrease in brain temperature and VB in the NREM phase [44, 58], as well as an increase in CO_2_ and acidity in the ICF [4, 59] initiate a switch from the GS1 mode to the GS2 mode (REM sleep).

GS2 is characterized by rapid eye movements, dreams, and, unlike GS1, a sharp increase in cerebral blood flow with expansion of the lumens of arterioles and capillaries [42, 43]. Considering the synesthesia of vision with almost all somatosensory systems [17, 60], we cannot exclude the convergence of the nervous systems of thermoreceptors and oculomotor muscles in the mechanism of synchronization of their activity [41]. Rapid eye movement enhances the communication of thermoreceptors of the cornea and cells of the ganglion layer of the retina through nerve connections with the thalamus, hypothalamus and midbrain substance. An increase in blood flow intensifies the water exchange necessary to wash out toxins from the parenchyma into the venules [42, 43, 47]. The expansion of the lumens of arterioles and capillaries enhances the polarization effects of the wall plasma layer in the electrophysics of synaptic plasticity [30]. The duration of GS2 is apparently determined by the time of depletion of the glucose supply in the extraocular muscles and the achievement of the threshold lactate value in them [42].

In the signaling systems of GS1 and GS2 a key role is played by the thermodynamics of the hydration shells of protein domains in aquaporin channels (AQP4) [30] and in voltage-dependent ion channels such as TRPM8 and TRPV1. Their physiological specialization is manifested in a significant difference in the duration and E_A_ of the NREM and REM phases of mice (Fig. 2). The effective E_A_=110 kJ/mol response, which determines the duration of the NREM phase (Fig. 2D), practically coincides with the E_A_=112 kJ/mol electrical stimulation of rat retinal ganglion cells *in vitro* [62]. This value follows from the F_A_ approximation of the TD threshold current of ganglion cells: 24.4 °C – 239 μA, 29.8 °C – 101 μA, 33.8 °C – 60 μA (R^2^=0.999) [62]. On the other hand, for the REM phase, the effective E_A_ = 270 kJ/mol (Fig. 2B) turns out to be of the same order of magnitude as the E_A_ of the TRPM8 threshold currents (Fig. 3). These results are consistent with the classification of eye thermoreceptors as regulators of sleep metabolism, and ganglion cells as activators of SCN – the pacemaker of the circadian rhythm. In mice, using iontophoresis, it was established that during sleep and anesthesia, the rate of diffusion of the tetramethylammonium cation in ICF increases by 60% [6, 45, 63]. It is known [64] that basal T in mice decreases by ∼14 °C 30 min after induction of anesthesia. Taking this into account, it can be assumed that in the T_w_–T_S_ range, the decay of HBs clusters in bulk water [30] initiates a gel-sol transition in ICF, as a result of which D_w_ and ion mobility increase by 60%. At the same time, an increase in the circulation of ICF in the GS1 mode will lead to an increase in the influx of CSF into the parenchyma from the subarachnoid space along the Virchow–Robin canals, which will widen due to the narrowing of the lumens of blood vessels under the influence of serotonin and norepinephrine produced by the pineal gland.

**Fig. 2.**
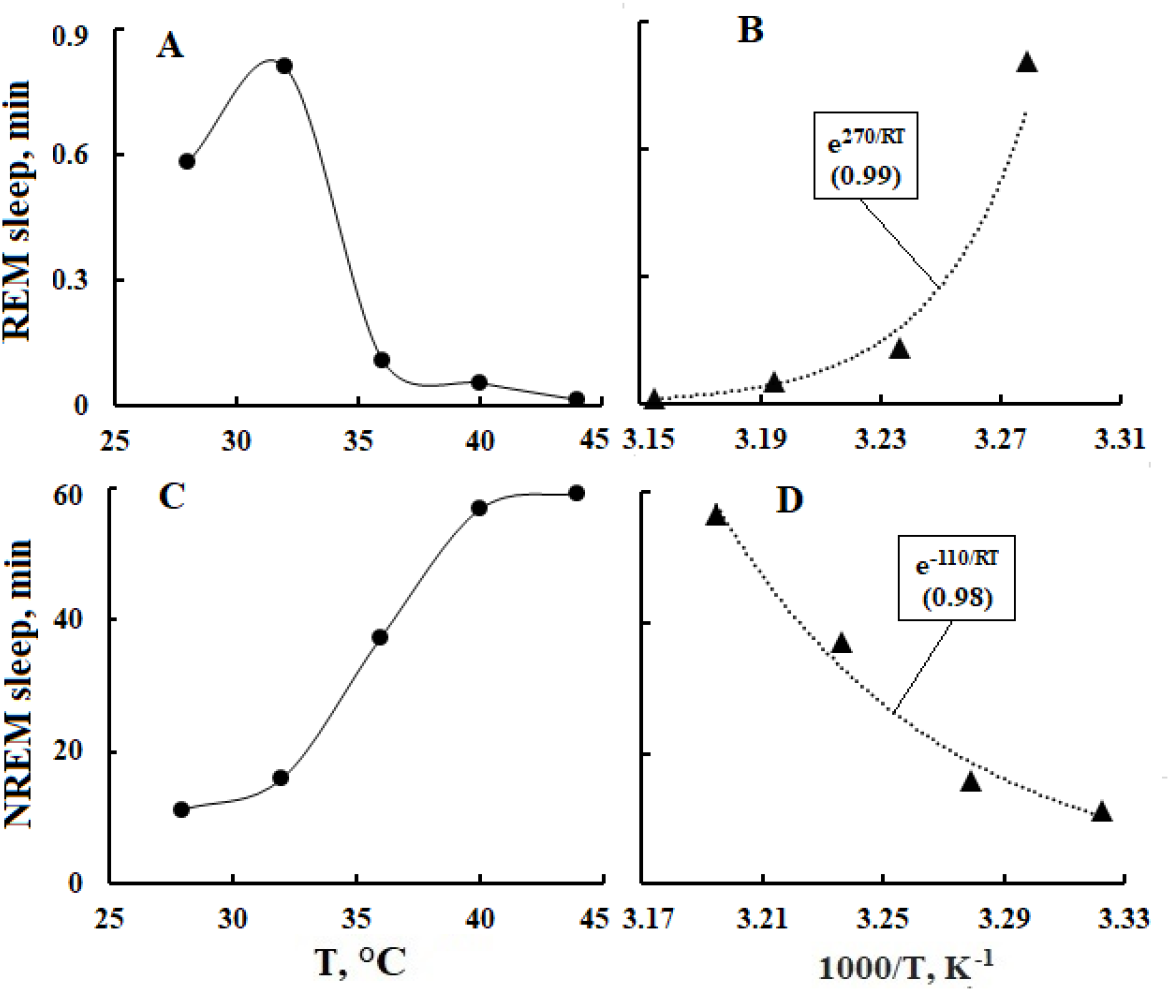
Points - dependence of the duration of sleep phases REM (**A, B**) and NREM (**C, D**) of mice on T (**A, C**) and 1/T (**B, D**) in the period of 23.00-01.00 hours; lines are envelopes (**A, B**) and F_A_ approximations (**B, D**). The number in the exponent in the box is equal to E_A_. Initial data from [61].

**Fig. 3.**
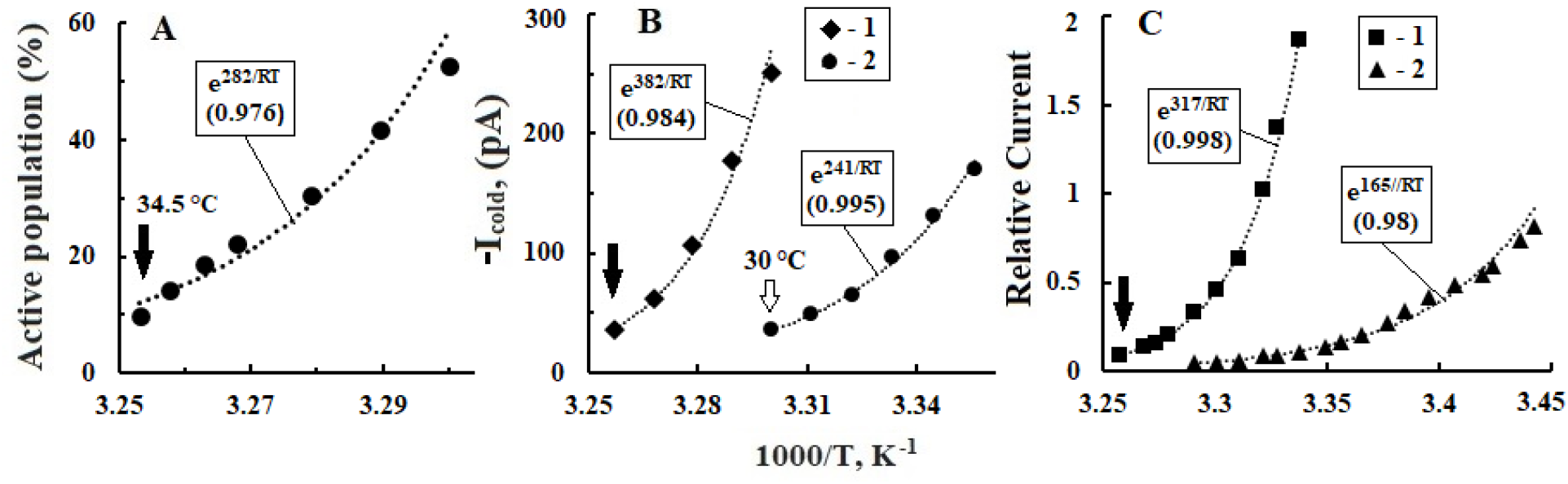
Points – dependences on 1/T of the bath. **A**, Cumulative distribution of active trigeminal neurons with TRPM8 channels. **B**, current in the TRPM8 channels of the trigeminal nerve (**1**) and the effect of on it of the addition of 1 μM BCTC (**2**). C, relative current in TRPM8 channels (**2**) at -60 mV and influence on it of 30 μM menthol (**1**). The arrows mark the threshold cold T. The lines are F_A_ approximations. Initial data **A** from [71], **B** from [15], **C** from [72].

### 3.3. Thermodynamics of thermo- and photoreceptors of the eye

During sleep, T_S_ = 36.5 °C have ICF, CSF of the third and lateral ventricles of the brain, as well as CSF of the subarachnoid space above the cortex [6, 45, 46, 65]. The temperature of the SCN and pineal gland will be close to T_S_, since they border the third ventricle of the brain [19, 66]. With external T=24±1.1 °C, the temperature under the tongue is 36.6±0.5 °C [1]. On the surface of the cornea, due to heat exchange with the external environment, T=34.7±1.1 °C is established, almost equal to T_w_ and the threshold T of the TRPM8 cold channel [15]. The VB temperature is 33.9±0.4 °C [1, 67], and the T of the retinal ganglion layer ranges from 34.8÷35.2 °C [67].

In the sleep state, T of the cornea and VB are lower than T_w_, therefore, the thermodynamics of eyeball fluids is dominated by clustering processes of HBs of bulk water [30]. Rearrangements of the hydration shells of proteins in TRPM8 channels and in VB fibrils at T<T_w_ contribute to an increase in the throughput of the former and initiate the processes of aggregation and crystallization of the latter [32, 68]. Indeed, the E_A_ values of TRPM8 operation (Fig. 2B, Fig. 3A) and the generation of ion currents (Fig. 3B, Fig. 3C and Fig. 4A) are comparable with the E_A_ of the processes of aggregation (∼50-70 kJ/mol) and crystallization (∼400 kJ/ mol) alpha helices in model solutions of high concentration hemoglobin’s [30].

**Fig. 4.**
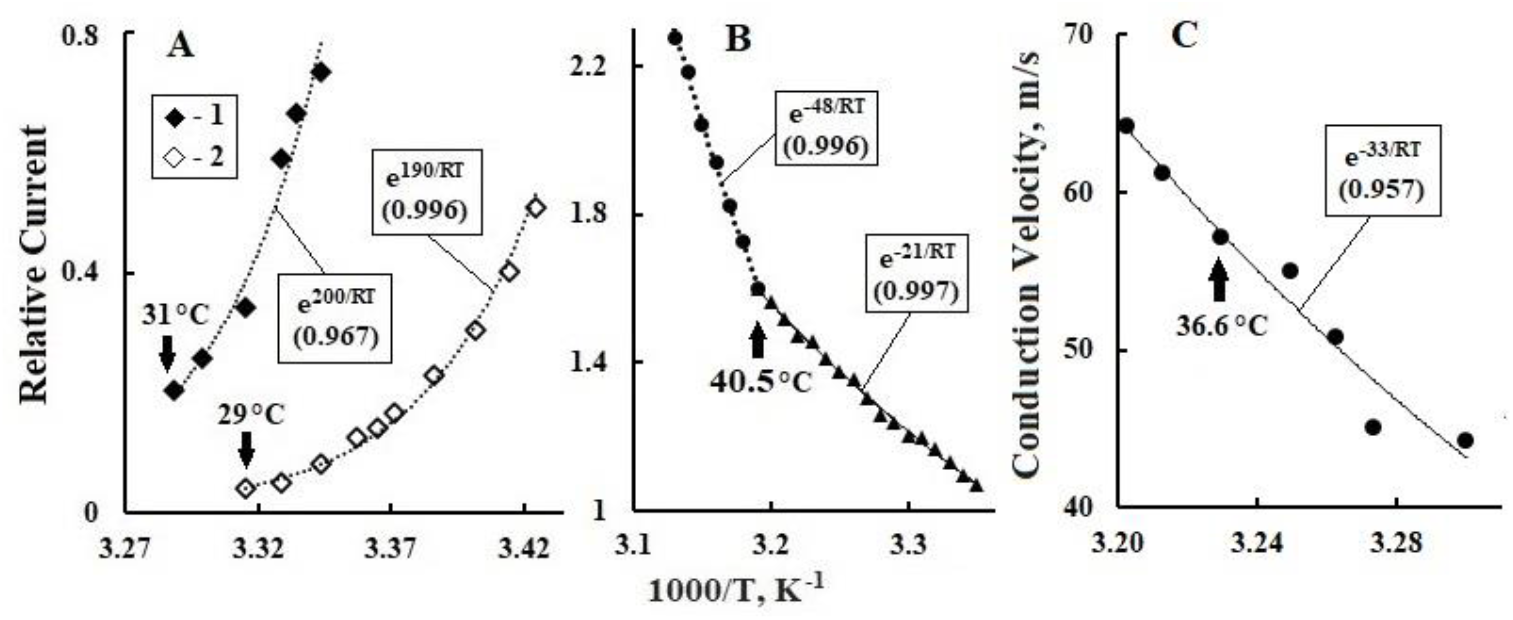
Dependences on 1/T of relative currents in various ion channels (points). The lines are F_A_ approximations. **A**, the TRPM8 channel of whole cells at a transmembrane potential of +100 (**1**) and -80 (**2**) mV, the arrows indicate the threshold cold T. **B**, the TRPV1 channel, the arrow marks the temperature of maximum sensitivity to heat. **C**, speed of action potential propagation between nodes of Ranvier of the myelinated fiber. Initial dependences on T: **A** from [72], **B** from [75], **C** from [78].

The influence of the dynamics of hydration shells on the E_A_ of TRPM8 currents is evidenced by a doubling of E_A_ when a specific TRPM8 activator, menthol, is added to the bath solution [16, 68, 69] (Fig. 3C) and a 1.5-fold decrease in the E_A_ current when the cold receptor blocker BCTC is added [15] (Fig. 3B). The OH group of menthol and the hydrophilic centers of BCTC, when interacting with the protein domains of the TRPM8 channels and the channels of the heat and pain receptors TRPV1, initiate charge redistribution in them between amino acids and cations [70-72]. In this case, local and transmembrane potentials arise [16, 73, 74], which control the mechanisms of opening and closing channels and ensure the passage of cations through them into or out of the neuron [15, 16]. The linear dependence of E_A_ currents TRPM8 on the potential difference follows from a comparison of TDs currents in Fig. 3C and Fig. 4A.

Corneal thermoreceptors with TRPV1 channels have maximum sensitivity at T∼40 °C [75], close to the pain threshold of 42 °C [17, 60, 76]. In the range from 25 to ∼40 °C, the E_A_ value of the ion current in the TRPV1 channel and in the bath buffer solution is 21 kJ/mol (Fig. 4B) and 17 kJ/mol [77], respectively. These E_A_ values correlate with the E_A_ of rotational-translational diffusion of water (τ_D_, D_w_) and electrolytes (T_1_) in the corresponding T ranges [30]. The increase in the E_A_ ion current in TRPV1 at T>40 °C to 48 kJ/mol is apparently due to the specific molecular mechanism of the TRPV1 response to T above the pain threshold [17, 75]. The average E_A_ current value in TRPV1 in the range of 33-42 °C is ∼34 kJ/mol, which coincides with the E_A_ (33 kJ/mol) TD of the rate of impulse transmission by the saltatory mechanism in the myelinated fiber (Fig. 4C).

This consistency of E_A_ values confirms the dependence of the kinetics of the saltatory mechanism on the efficiency of the voltage-dependent K^+^ channels in the nodes of Ranvier [33]. The saltatory mechanism still operates on fibers with the myelin sheath removed at T<36.5°C, but is blocked at T>36.5°C [78]. The blocking can be associated with the destruction at T>36.5 °C of the HBs structure in the axoplasm, which is necessary for the generation and propagation of polarization waves between the nodes, activating K^+^ channels in them [33]. It also follows that in sleep at brain T∼35÷36.5 °C, the speed of action potential transmission to the SCN along ipRGC axons, which do not have myelin sheaths within the retina [79], will be about 50 m/s (Fig. 4C). Let us assume that activation of the retina by light upon awakening is associated with an increase in T axons of ganglion cells outside the retina to T_b_ (∼37 °C) and in the case of ipRGC this leads to blocking their connection with the SCN [78]. In this way, changes in external T can initiate the triggering of the mechanism for switching brain homeostasis from daytime to nighttime mode [6, 34].

## 4. Conclusion

The systematic analysis of the known temperature dependences of the kinetic characteristics of the signaling and trophic functions of the brain carried out in the work showed that the electrical and dynamic properties of water play a key role in their molecular mechanisms. This is confirmed by correlations between activation energies of the temperature dependences of hydrogen bonds in the hydration shells of proteins and the dynamic parameters of physiological fluids. The synergism of thermal librational vibrations of water molecules and exothermic proton hopping determines the thermodynamics of fluids in the human eyeball and brain in the range of 33-40 °C. During the process of phylogenesis in the physiology of terrestrial mammals, a mechanism of adaptation to the circadian rhythm has developed, taking into account the specifics of the lifestyle and the physical characteristics of the habitat. This mechanism is based on the integration of the visual system with almost all somatosensory and the localization in the eyeball of light and cold receptors, which are responsible for switching the functions of the pineal gland and suprachiasmatic nucleus of the hypothalamus from day to night mode. This is confirmed by correlations between the duration of sleep-in mammals and the density of neurons in the neocortex, the diameter of the eyeball, and the mass of the pineal gland. In addition, there is a strict consistency between the nighttime temperature of the eyeball and the threshold for triggering the TRPM8 channel of the corneal cold receptor, as well as the daytime temperature of the brain with the threshold for blocking the communication channel of the retinal ganglion cells ipRGC with the suprachiasmatic nucleus. The nerve connections of the eyeball with the brain are able to divide the nocturnal metabolism of the brain into two phases of sleep – NREM and REM, which correspond to two modes of the glymphatic system of the brain – electrochemical and dynamic. The first mode is characterized by relaxation processes of synaptic plasticity and chemical neutralization of toxins with the participation of water and melatonin. Activation of the extraocular muscles and a sharp increase in cerebral blood flow in the second mode intensify water exchange in the parenchyma and the leaching of toxins into the venous system of the brain. From the results of the work, it follows that classical electrophysics and thermodynamics of water underlie the neurophysiology of the basic functions of the brain, which are characteristic of all mammals, including humans. It can be hoped that deepening the knowledge of the properties of water to the level of subquantum physics will make it possible to study the nature of the physical uniqueness of the human mind [33, 40, 80, 81].

## Availability of data and materials

The initial data used for the figures and tables of this study are available in the articles, links to which are given in the captions to figures.

